# Innate immune cell-intrinsic ketogenesis is dispensable for organismal metabolism and age-related inflammation

**DOI:** 10.1101/2022.10.14.512312

**Authors:** Emily L. Goldberg, Anudari Letian, Tamara Dlugos, Claire Leveau, Vishwa Deep Dixit

## Abstract

Aging is accompanied by chronic low-grade inflammation, but the mechanisms that allow this to persist are not well understood. Ketone bodies are alternative fuels produced when glucose is limited and improve indicators of healthspan in aging mouse models. Moreover, the most abundant ketone body, β-hydroxybutyrate (BHB), inhibits the NLRP3 inflammasome in myeloid cells, a key potentiator of age-related inflammation. Given that myeloid cells express ketogenic machinery, we hypothesized this pathway may serve as a metabolic checkpoint of inflammation. To test this hypothesis, we conditionally ablated ketogenesis by disrupting expression of the terminal enzyme required for ketogenesis, 3-Hydroxy-3-Methylglutaryl-CoA Lyase (HMGCL). By deleting HMGCL in the liver, we validated the functional targeting and establish that the liver is the only organ that can produce the life-sustaining quantities of ketone bodies required for survival during fasting or ketogenic diet feeding. Conditional ablation of HMGCL in neutrophils and macrophages had modest effects on body weight and glucose tolerance in agin, but worsened glucose homeostasis in myeloid cell specific Hmgcl deficient mice fed a high-fat diet. Our results suggest that during aging, liver derived circulating ketone bodies might be more important for deactivating NLRP3 inflammasome and controlling organismal metabolism.

## Introduction

Mammals have evolved to prioritize glucose for energy. A complex, carefully regulated system has developed to control glucose availability and utilization for every cell in the body. However, during periods of starvation or limited glucose availability, mammals break down fat, leading to the production of ketone bodies, to supply energetic demand (Cahill, 2006). Ketone bodies are short chain fatty acids that fuel cellular ATP production through their ability to enter the TCA cycle. Thus, ketone bodies are often considered alternative metabolic fuels. Notably, many metabolic interventions that induce ketogenesis also extend lifespan in model organisms (Edwards et al., 2014; Fang et al., 2013; Kuhla et al., 2014; Sengupta et al., 2010). Moreover, ketogenic diets (KD) improve markers of healthspan in old mice (Newman et al., 2017; Roberts et al., 2017). Collectively, these studies underscore an important role for ketone bodies in aging and healthspan.

The metabolism of ketone bodies has been expertly reviewed previously (Puchalska and Crawford, 2021). Free fatty acids liberated from adipose tissue through lipolysis are broken down through β-oxidation in the liver, leading to the production of acetyl-CoA. As the concentration of acetyl-CoA increases in hepatocyte mitochondria it is converted to ketone bodies through a series of enzymatic reactions. The final step in ketogenesis is catalyzed by the enzyme 3-Hydroxy-3-methylglutaryl-CoA lyase (HMGCL, encoded by the gene *Hmgcl*) to form the ketone body acetoacetate, which can then be converted to the other ketone bodies β-hydroxybutyrate (BHB) and acetone. Hepatocytes do not express the enzyme required for ketolysis Succinyl-CoA:3-Ketoacid-CoA Transferase (SCOT, encoded by the gene *Oxct1*), and this preserves ketone bodies for extrahepatic tissues like the brain, heart, and skeletal muscle (Fukao et al., 1997).

In addition to this classical regulation of ketogenesis, recent evidence shows non-hepatic sources of ketone bodies impact a variety of organ systems. In adipose tissue, beige PRDM+ adipocytes secrete BHB that is oxidized by adipocyte precursors to preserve adipogenic differentiation and limit fibrotic lineage skewing (Lecoutre et al., 2022; Wang et al., 2019). BHB is produced by small intestine Lgr5+ stem cells and this is important for maintaining their stemness within crypts (Cheng et al., 2019; Stine et al., 2019). Local ketone production has been reported in CD8 T cells and implicated to regulate their memory response (Zhang et al., 2020a). Renal epithelial cells produce BHB to mediate protective effects of nicotinamide (NAM) (Tran et al., 2016). The failing heart also increases ketone body consumption (Horton et al., 2019; McCommis et al., 2020; Zhang et al., 2020b). Finally, macrophages can oxidize acetoacetate depending on their inflammatory state, and this is important for protecting against liver fibrosis (Puchalska et al., 2018a; Puchalska et al., 2018b). Collectively, these studies emphasize that ketone bodies may have autocrine/paracrine functions and have broad physiological importance.

Ketone bodies are also pleiotropic signaling molecules. BHB acts as a histone deacetylase inhibitor to control gene expression (Shimazu et al., 2013). Similar to other short chain fatty acids, BHB can covalently bind lysine residues on histones and other proteins, although the importance of this post-translational modification is not well understood (Koronowski et al., 2021; Terranova et al., 2021; Xie et al., 2016; Zhang et al., 2020a). In addition, we previously showed that BHB inhibits NLRP3 inflammasome activation in macrophages (Youm et al., 2015) and neutrophils (Goldberg et al., 2017). Persistent low-grade inflammation is believed to underlie many diseases of aging (Franceschi and Campisi, 2014) and we have previously shown that the NLRP3 inflammasome is a key driver of age-related and obesity-driven inflammation (Vandanmagsar et al., 2011; Youm et al., 2013). Based on the broad actions of BHB, we hypothesized that ketone bodies might be an important regulatory checkpoint for chronic inflammation in aging.

Ketone bodies impact a wide range of immune functions (Ang et al., 2020; Goldberg et al., 2017; Goldberg et al., 2019; Karagiannis et al., 2022; Liu et al., 2022; Luda et al., 2022; Puchalska et al., 2018b; Ryu et al., 2021; Thio et al., 2022; Youm et al., 2015; Zhang et al., 2020a). While several studies have investigated the fate of extracellular ketone bodies in immune function, less is known about immune cell-intrinsic ketogenesis. To test the importance of ketone body production in macrophages and neutrophils, we developed a novel mouse model by targeting *Hmgcl* (*Hmgcl^fl/fl^*) to conditionally ablate ketone body synthesis in specific cell types. This strategy allowed us to focus exclusively on ketone body production, in contrast to pre-existing models targeting the upstream rate-limiting enzyme HMGCS2 (Cheng et al., 2019), or the downstream BDH1 that interconverts acetoacetate to BHB (Horton et al., 2019). By crossing these mice to liver-specific Albumin-Cre (Hmgcl^Alb-Cre^) we show that despite the presence of non-hepatic ketogenesis, the liver is the only organ that can produce enough ketone bodies to support survival under ketogenic conditions. We also find that neutrophil (using S100a8-Cre, Hmgcl^S100a8-Cre^)-intrinsic ketogenesis does not regulate age-related metabolic health. In addition, using LysM-Cre to ablate ketogenesis in all myeloid cells (Hmgcl^LysM-Cre^), we find only modest impacts on age- and obesity-induced metabolic dysregulation. These data suggest that innate immune inflammation is controlled by extracellular ketone bodies, and that innate immune-intrinsic ketogenesis does not regulate age-related inflammation and metabolic health defects in aging.

## Results

To test the role of ketogenesis within innate immune cells, we first generated a mouse model containing a loxP-flanked region of exon 2 within the *Hmgcl* gene and verified homozygosity in genomic DNA (Fig 1A, B). To validate functional gene targeting, we first crossed these Hmgcl^fl/fl^ mice with the liver-specific Albumin-Cre (Hmgcl^Alb-Cre^) and confirmed protein deletion (Fig 1C). In contrast to whole-body HMGCL deficiency that is embryonic lethal (Wang et al., 1998), liver specific Hmgcl deficient mice fed a normal chow diet were viable. When fed a ketogenic diet (KD), Hmgcl^Alb-Cre^ mice failed to increase circulating blood BHB concentration (Fig 1D) and Cre+ mice had lower blood glucose (Fig 1E). Our data agree with a prior study in another Hmgcl^fl/fl^ model that was published while we were developing our mice (Gauthier et al., 2013). Interestingly, when fed KD, the mice also fail to induce lysine β-hydroxybutyrlation (referred to as Kbhb) (Fig 1C), a newly described post-translational modification by BHB (Xie et al., 2016). Concomitant with their inability to induce hepatic ketogenesis, Hmgcl^Alb-Cre^ mice failed to maintain body weight during KD feeding (Fig 1F), primarily due to increased adipose tissue lipolysis (Fig 1G). Notably, the weight-loss phenotype could be rescued if we cycled mice off of KD every 24 hours in exchange for standard chow (Fig 1H) demonstrating the specificity of HMGCL-mediated hepatic ketogenesis in maintaining the body weight. In addition to KD, Hmgcl^Alb-Cre^ mice also fail to induce ketogenesis in response to fasting (Fig 1I) and they have lower blood glucose levels under fasting conditions (Fig 1J). All together, these data functionally validate HMGCL ablation in this new mouse model and demonstrate the liver is the only organ that can supply enough ketone body production under ketogenic conditions, and that no other tissues can compensate for hepatic ketogenesis to meet whole-body energy requirements during ketogenic diet feeding or fasting.

**Figure 1.**
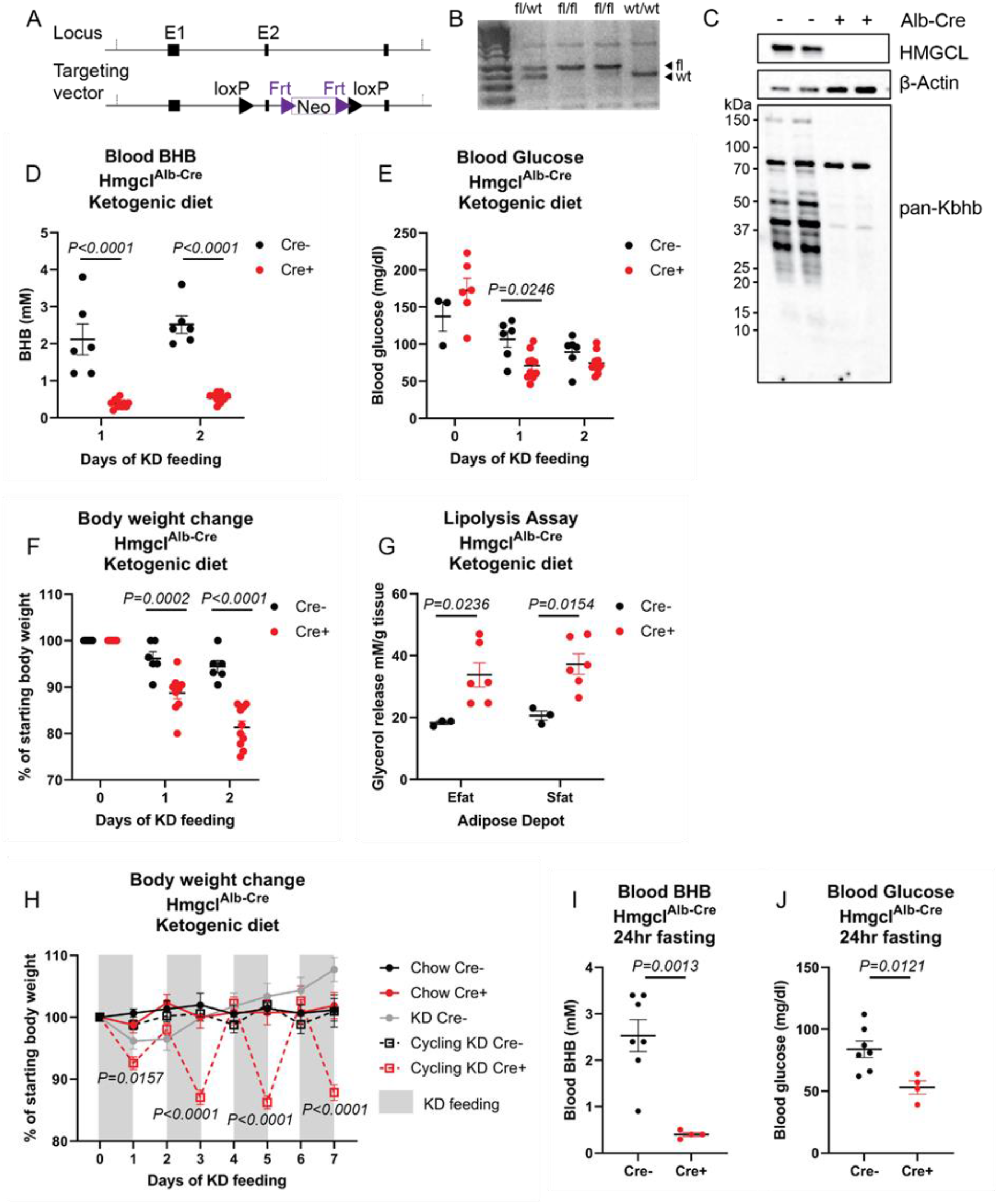
Development and validation of a novel mouse model to conditionally ablate ketogenesis. A mouse was generated to study cell-specific ketogenesis by targeting expression of *Hmgcl*. (A) Targeting vector design to introduce loxP sites to allow Cre-mediated excision of exon 2 within the *Hmgcl* gene. (B) Representative genotyping gel. Liver-specific *Hmgcl* ablated mice were fed a ketogenic diet. (C) HMGCL, Actin, and Kbhb were assessed in livers of Cre+ and Cre-Hmgcl^Alb-Cre^ mice. Blood (D) BHB, (E) blood glucose, and (F) body weights were measured each morning. (G) Glycerol release from Efat and Sfat was measured after 48hrs of KD feeding. For D-G, all statistical differences were calculated by 2-way ANOVA to compare genotypes at each time point. (H) Body weights were measured each morning during 1 week of cycling KD feeding. Statistical differences between Cre+ and Cre-Hmgcl^Alb-Cre^ littermates were calculated by 2-way ANOVA. Data are represented as mean±SEM. (I) Blood BHB and (J) blood glucose levels were measured

Based on our prior findings that neutrophil NLRP3 inflammasome activation can be regulated by BHB and the unexpected expression of ketogenic enzymes in these short-lived glycolytic immune cells (Fig 2), we tested the importance of neutrophil-intrinsic ketogenesis by crossing the Hmgcl^fl/fl^ mice to the neutrophil-specific S100a8-Cre (Hmgcl^S100a8-Cre^). Cre specificity was assessed by crossing to the mTmG reporter mouse, mTmG^S100a8-Cre^, (Fig 3A, B). We stimulated NLRP3 inflammasome activation in isolated bone marrow neutrophils and found that HMGCL does not regulate NLRP3-dependent IL-1β secretion (Fig 3C). The immune phenotyping of adult male and female mice revealed that HMGCL ablation in neutrophils does not impact peripheral neutrophil abundance, nor does it have indirect effects on other lymphoid populations (Fig 3D-G). These data confirm efficient deletion of HMGCL using S100a8-Cre and this does not impact overall neutrophil abundance in the periphery.

**Figure 2.**
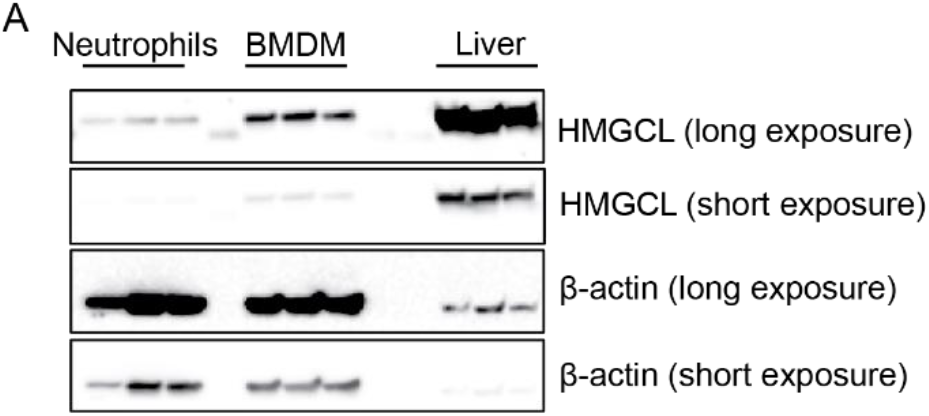
Comparison of HMGCL expression between liver and myeloid cells. (A) HMGCL protein expression was compared in neutrophils, bone marrow-derived macrophages (BMDM), and whole liver tissue. Short and long exposures are provided as indicated. Each lane is an individual mouse.

**Figure 3.**
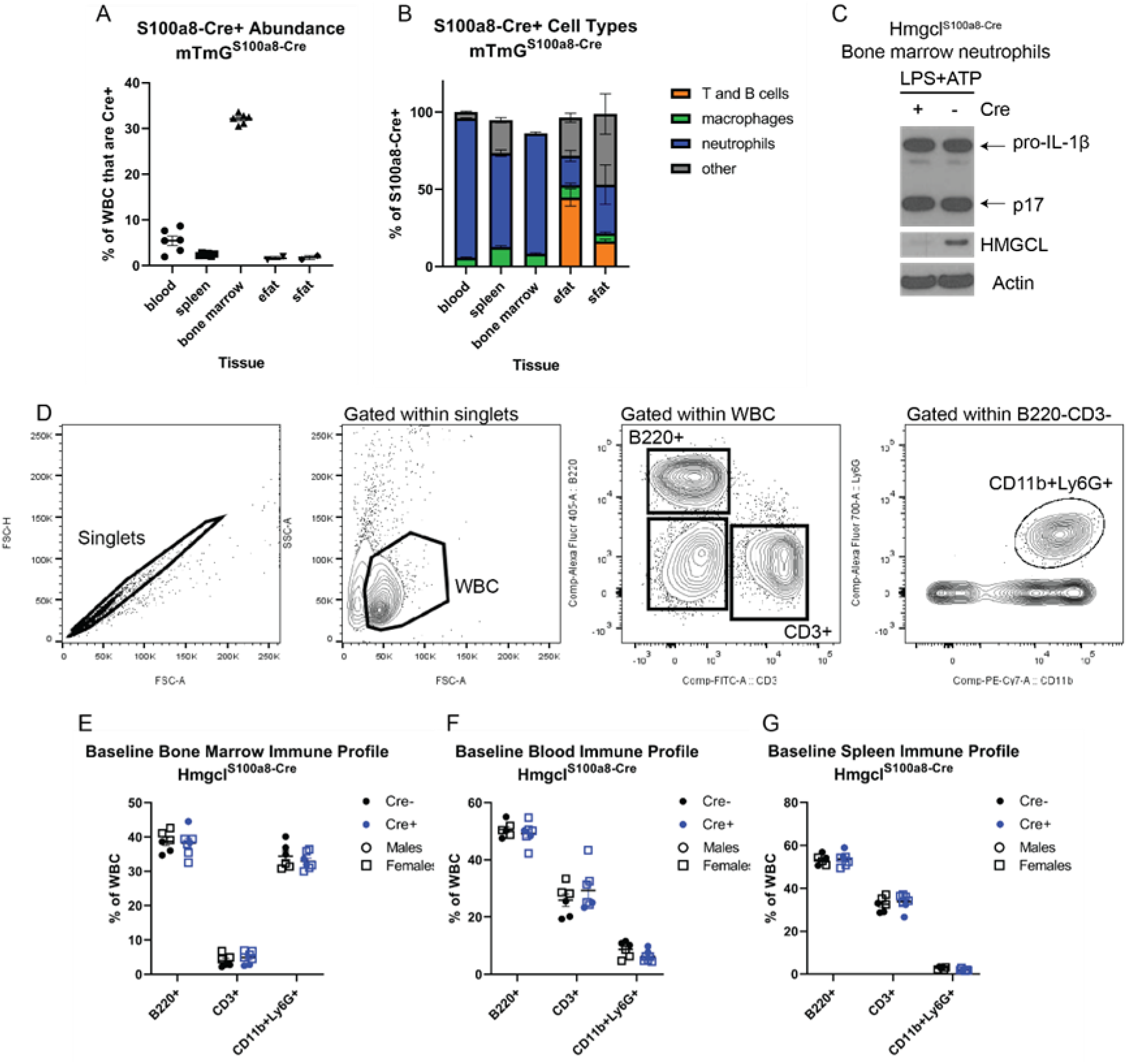
Validation and baseline characterization of HMGCL ablation in neutrophils. For (A-B) S100a8-Cre^+^ mice were crossed to mTmG reporter mice to verify neutrophil-specificity. (A) The total abundance of Cre+ (mGFP+) cells in each tissue were measured by flow cytometry. (B) The cell lineages within the Cre+ cells of each tissue were determined by flow cytometry. For both graphs, each symbol represents an individual mouse and all data are expressed as mean±SEM. (C) Representative western blot of isolated bone marrow neutrophils from Hmgcl^S100a8-Cre^ mice after NLRP3 inflammasome activation with LP≡+ATP. (D) Representative flow cytometry gating strategy to define B220+ B cells, CD3+ T cells, and CD11b+Ly6G+ neutrophils. For (E-G) Hmgcl^fl/fl^S100a8-Cre^+/-^ mice were compared to Hmgcl^fl/fl^ littermates to test the role of ketogenesis within neutrophils. Baseline B220+ B cells, CD3+ T cells, and neutrophils were quantified in (E) bone marrow, (F) blood, and (G) spleen. Male (circles) and female (squares) Cre+ (black) and Cre-(blue) littermates were combined for analysis and each symbol represents an individual mouse.

Upon aging, male Hmgcl^S100a8-Cre^ mice, showed modest differences in body weights (Fig 4A). Likewise, neutrophil-specific deletion of *Hmgcl* also modestly protected 15-20 month-old male mice from age-related glucose intolerance compared to male Cre-negative littermate controls (Fig 4B). To test if HMGCL ablation altered acute inflammatory responses, we analyzed physiological responses to intraperitoneal injection with gram-negative bacterial cell wall component lipopolysaccharide (LPS) that activates TLR4 signaling. While we measured expected changes in blood glucose and body temperature in LPS-challenged mice, there was no difference between Cre-positive and Cre-negative mice (Fig 4C-E). Collectively, these data suggest that neutrophil-intrinsic HMGCL expression, and hence neutrophil-intrinsic ketogenesis, is not a major regulator of inflammation during aging.

**Figure 4.**
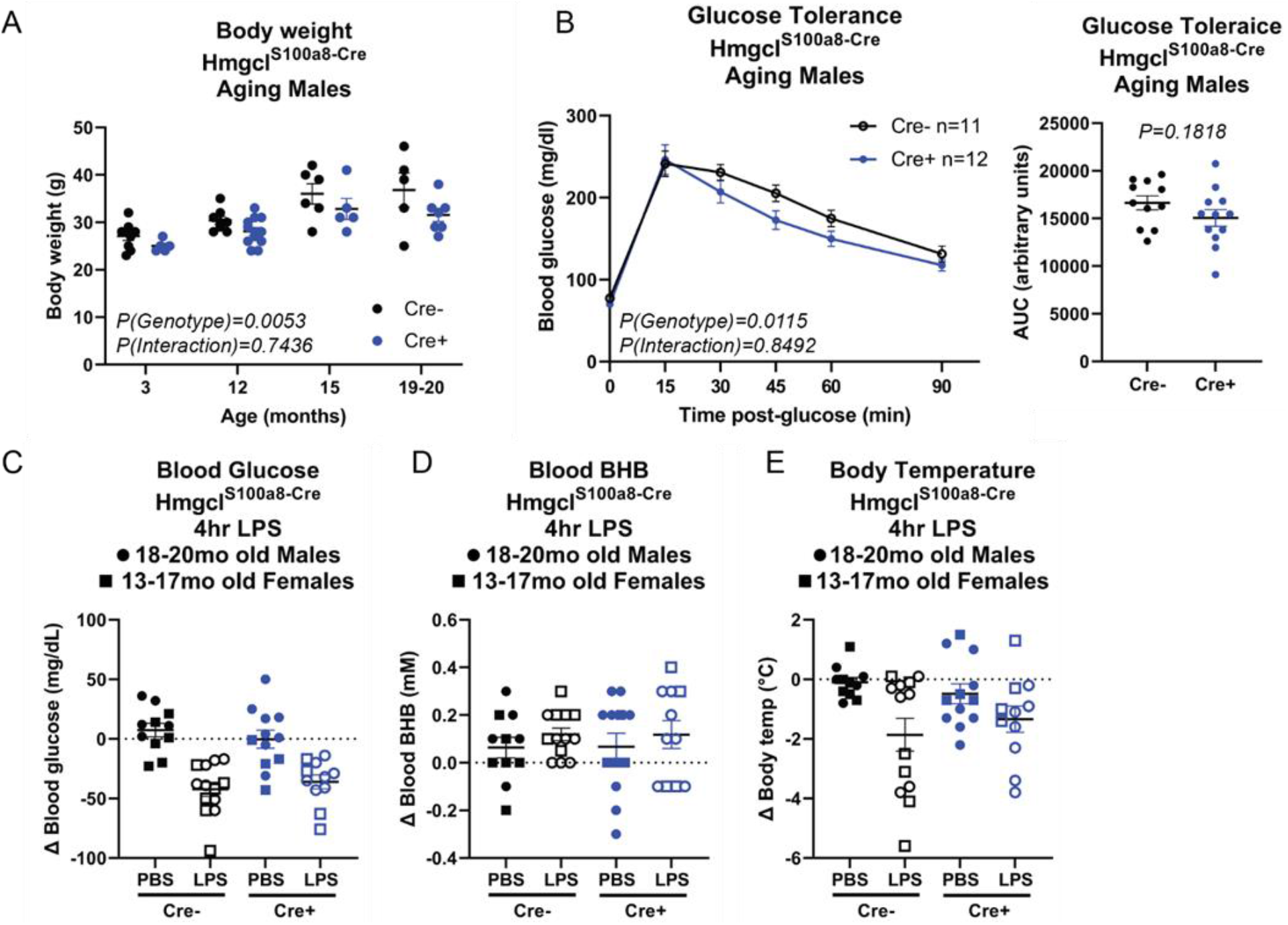
Neutrophil-specific *Hmgcl* ablation does not impact age-related inflammation. Hmgcl^S100a8-Cre^ male mice were aged to at least 18 months old. (A) Body weights of independent cohorts at varying ages. Statistical differences were calculated by 2-way ANOVA. (B) Glucose tolerance test of males aged 15-18 months old. Area under the curve was quantified for each animal (right side panel). For (C-E) 18-20 month-old males (circles) and 13-17 month-old females (squares) were injected with LPS or PBS control and (F) blood glucose, (G) blood BHB, and (H) body temperature were measured 4 hours later to assess the physiological response to acute inflammation. Statistical differences were calculated by paired 2-way ANOVA. For all graphs, each symbol represents an individual mouse and all data are expressed as mean±SEM.

Because macrophage function can be regulated by ketone bodies (Puchalska et al., 2018b; Youm et al., 2015), we broadened our scope by assessing the role of HMGCL in all myeloid-lineage cells. For these experiments, we crossed Hmgcl^fl/fl^ mice with mice expressing the generic myeloid LysM-Cre driver (Hmgcl^LysM-Cre^) to ablate HMGCL in all myeloid cells *in vivo*. After one week of KD feeding, Cre-positive and Cre-negative adult littermates had similar blood BHB levels (Fig 5A), confirming that myeloid cells do not contribute to whole-body circulating BHB. In contrast to the modest phenotype in aged Hmgcl^S100a8-Cre^ mice, Hmgcl^LysM-Cre^ mice had no differences in body weight, glucose tolerance, or fasting-induced weight loss in 18 month-old male Cre-positive and Cre-negative littermate controls (Fig 5B-D).

**Figure 5.**
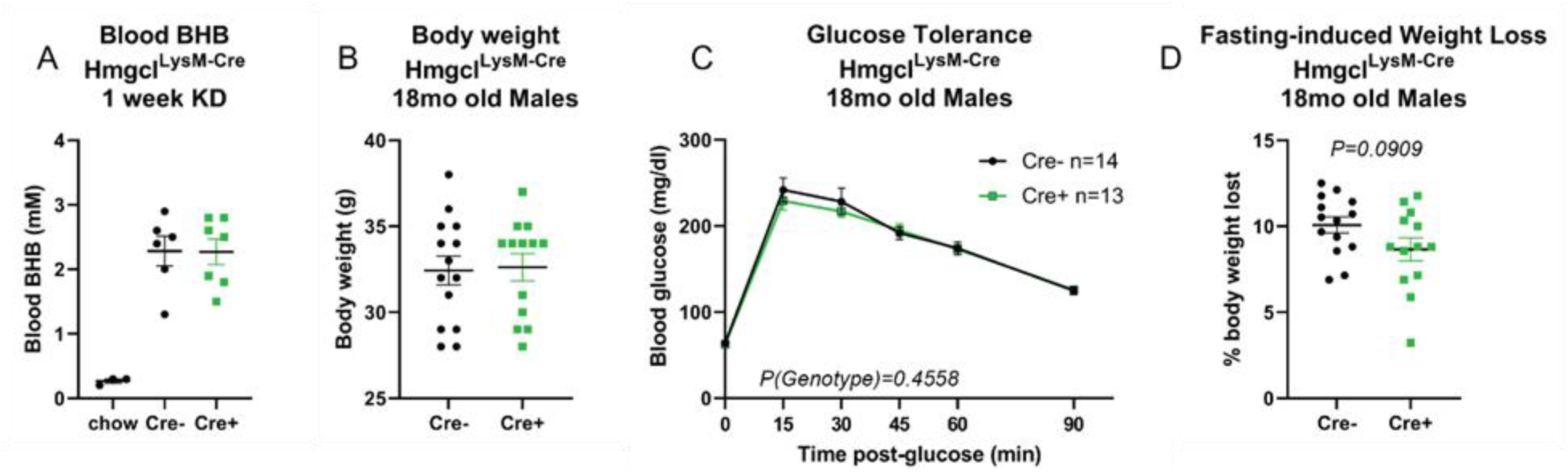
Myeloid-specific *Hmgcl* expression does not regulate metabolic health during aging. Hmgcl^LysM-Cre^ male mice were aged to 18 months-old and assessed for basic metabolic health parameters compared to their Cre-negative Hmgcl^fl/fl^ littermates. (A) Hmgcl^LysM-Cre^ were fed KD for 1 week to measure blood BHB levels. (B) Body weights, (C) glucose tolerance, and (D) 16-hour fasting-induced weight loss were measured. For all graphs, each symbol represents an individual mouse. For A, B, and C, statistical differences were calculated by unpaired student’s t-test. For C, paired 2-way ANOVA was used to compare statistical differences between genotypes and all data are expressed as mean±SEM.

Next, we tested if myeloid cell-intrinsic ketogenesis regulates obesity-induced inflammation. Male Hmgcl^LysM-Cre^ mice were fed a high-fat diet for 12 weeks to induce obesity. Despite no differences in body weight or fasting blood glucose, Cre-positive mice had lower glucose tolerance compared to Cre-negative littermates (Fig 6A-C). However, both genotypes had similar prevalence of adipose tissue macrophages in their epididymal adipose tissue (Fig 6D). Likewise, we measured no difference in M1/M2 polarization in Hmgcl-sufficient and Hmgcl-deficient bone marrow-derived macrophages in vitro (Fig 6E-H). These data suggest the importance of ketogenesis in innate immune cells may depend on both cell type and physiological state.

**Figure 6.**
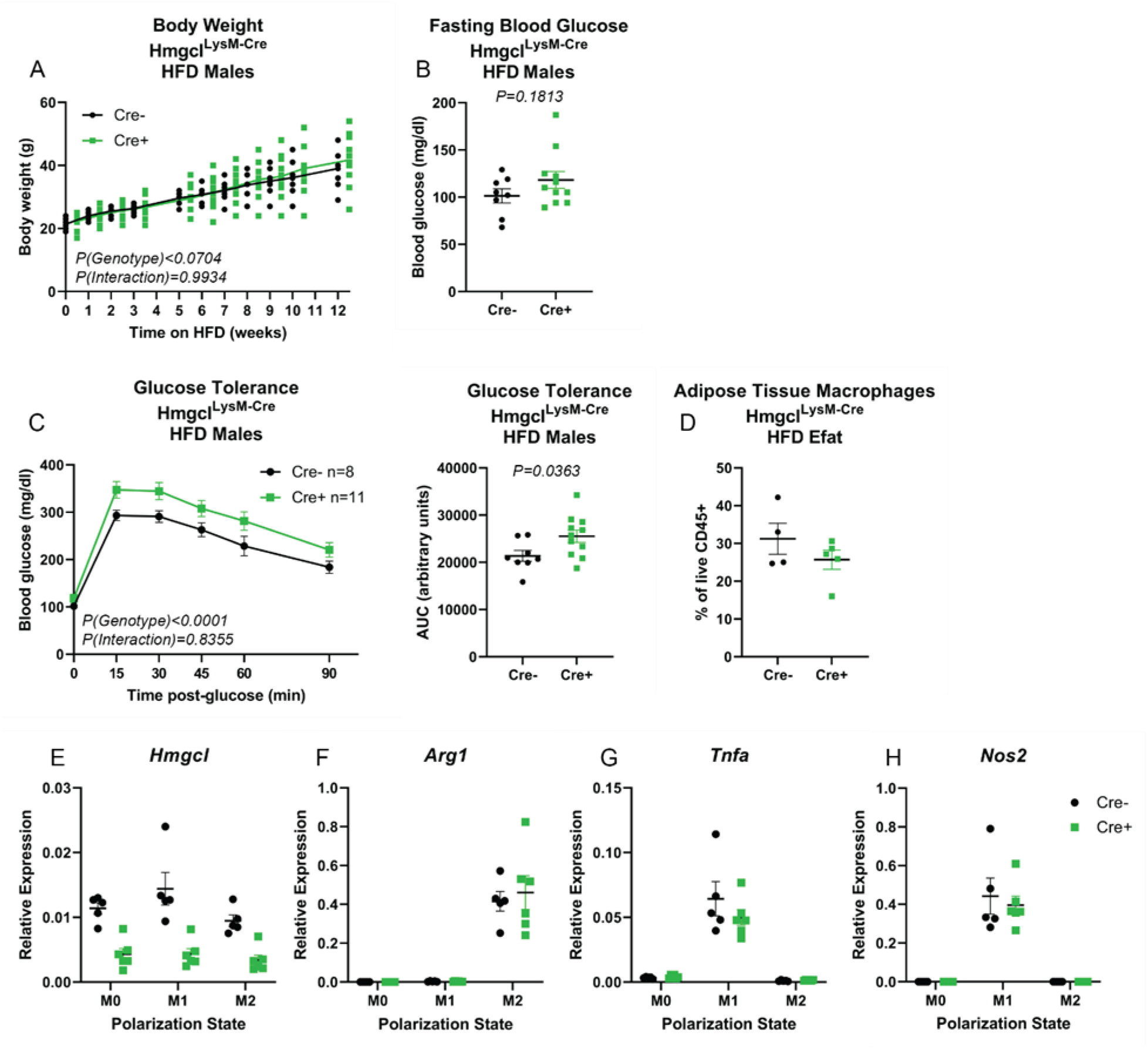
Myeloid-specific *Hmgcl* expression modestly impacts metabolic phenotypes in high-fat diet-induced obese mice. Male Hmgcl^LysM-Cre^ mice were fed a high-fat diet for 12 weeks to induce obesity. (A) Body weights were measured weekly throughout the experiment. (B) 16-hour fasting blood glucose and (C) glucose tolerance was measured. For (A, B) each symbol represents an individual mouse. Statistical differences were calculated by paired 2-way ANOVA and AUC differences were calculated by student’s t-test. (D) Abundance of adipose tissue macrophages was determined by flow cytometry. Bone marrow-derived macrophages from Hmgcl^LysM-Cre^ mice were used to measure M1/M2 macrophage polarization by analyzing (E) *Hmgcl*, (F) *Arg1*, (G) *Tnfa*, and (H) *Nos2* gene expression by RT-PCR. For D-H each symbol represents a sample that is pooled from n=2 mice. All data are expressed as mean±≡EM.

## Discussion

Prior studies have conditionally ablated ketogenesis in extrahepatic tissues by targeting the pathway rate-limiting enzyme HMGCS2 (Cheng et al., 2019; Venable et al., 2022). However, this enzyme catalyzes the production of β-hydroxy-β-methylglutaryl CoA (HMG-CoA) from acetoacetyl CoA. HMG-CoA is then converted to acetoacetate by HMGCL. Therefore, possible confounding effects of loss of HMG-CoA cannot be ruled out. Notably, HMG-CoA can also be produced from leucine catabolism, so our approach blocks ketogenesis from this pathway as well. To limit our focus exclusively on ketone body synthesis, we developed the *Hmgcf^fl/fl^* mouse to ablate the terminal enzyme leading to production of all three ketone bodies, acetoacetate, BHB, and acetone. We validated the functional targeting of *Hmgcl* disruption in the Hmgcl^Alb-Cre^ mice by showing these mice fail to upregulate ketogenesis in response to ketogenic diet and fasting. Our results in the Hmgcl^Alb-Cre^ mice also formally demonstrate that liver is the only organ that supports life-sustaining ketone body production during ketogenic diet feeding and fasting, and that all other combined sources of ketogenesis cannot compensate for the loss of hepatic ketogenesis.

The importance of ketone body production in non-hepatocytes is not completely understood. We and others have shown that immune cells are highly sensitive to ketone bodies through metabolic and non-metabolic mechanisms (Ang et al., 2020; Goldberg et al., 2017; Goldberg et al., 2019; Karagiannis et al., 2022; Liu et al., 2022; Luda et al., 2022; Puchalska et al., 2018b; Ryu et al., 2021; Thio et al., 2022; Youm et al., 2015; Zhang et al., 2020a). Moreover, the source of ketone bodies in immune modulation has not been defined and is likely different for each cell type and physiological condition. It is especially perplexing that short-lived neutrophils, with no obvious metabolic reliance on ketone bodies, would express ketogenic or ketolytic pathways. Given that innate immune cells express ketogenic enzymes and that their functions are impacted by both BHB and acetoacetate (Goldberg et al., 2017; Puchalska et al., 2018b; Youm et al., 2015), we designed this study to define the role of neutrophil- and macrophage-intrinsic ketogenesis in regulating inflammation and metabolic health during aging. Because both BHB and acetoacetate have anti-inflammatory roles in macrophages, we expected the deletion of HMGCL in these cells to cause elevated inflammation that would exaggerate metabolic dysfunction in old mice. Surprisingly, we found only modest effects of conditional HMGCL ablation on physiological indicators of metabolic health in aged mice. In aged male neutrophil-specific Hmgcl^S100a8-Cre^ mice, we observed reproducible but small differences in body weights during aging, and this translated to corresponding slight improvement in glucose tolerance. However, Hmgcl^S100a8-Cre^ mice showed no change to acute inflammatory challenge with LPS. In contrast, mice with broader disruption of ketogenesis in all myeloid-lineage cells (Hmgcl^LysM-Cre^) did not replicate the aging phenotypes seen in Hmgcl^S100a8-Cre^ mice. These data raise the possibility that ketogenesis in different innate immune cell types may have different competing effects on whole-body physiology. Our data also suggest that exogenous ketone bodies from other organs are important for controlling age-related neutrophil and macrophage inflammation. Unfortunately, due to the >20% weight loss, that necessitates euthanasia of Hmgcl^Alb-Cre^ mice under ketogenic conditions, we were unable to directly test this possibility.

The results from our study still do not explain why innate immune cells express ketogenic enzymes. This is particularly intriguing in neutrophils, which are short-lived and highly glycolytic, and therefore have no obvious metabolic requirement for ketone bodies. Moreover, the lack of *Bdh1* expression in macrophages and neutrophils limits their potential metabolic ketone body utilization to acetoacetate, although this does not preclude a non-metabolic role for exogenous BHB. We suspect that the role of ketone pathways in innate immune cells may be cell type- and disease-specific. We studied this in the context of aging and obesity due to our prior work linking NLRP3 inflammasome activation to metabolic inflammation in these conditions (Vandanmagsar et al., 2011; Youm et al., 2013), and our subsequent discovery that BHB inhibits the NLRP3 inflammasome (Goldberg et al., 2017; Youm et al., 2015). However, our *in vitro* data show that cell-intrinsic ketogenesis does not impact neutrophil NLRP3 activation or macrophage polarization. These data are in agreement with our prior work that acetoacetate does not inhibit NLRP3 inflammasome activation, and that these cells likely do not produce their own BHB. These data probably also explain, at least in part, the modest phenotypes we observed *in vivo*. Future studies using *in vivo* substrate tracing testing alternative sources of ketone bodies, and using additional disease models, should be carefully considered for testing the role of ketone pathways in immune cells.

## Materials and Methods

### Animals

All mice were housed in specific pathogen-free conditions under normal 12hr light/dark cycles. *Hmgcl*^fl/fl^ mice were generated by the Pennington Biomedical Research Center Transgenics Core. Genetic targeting to insert loxP sights into exon 2 of the *Hmgcl* allele was achieved by recombination of a PL253-loxP-frt-neo cassette in mouse embryonic stem cells. Upon confirmation of desired vector design and neomycin selection for initial enrichment of the targeted clone, we used these embryonic stem cells for injection into blastocysts for the generation of heterozygous Hmgcl-loxP floxed mice. These positive ES cells were used. Using polymerase chain reaction to genotype these neo-founder mice, we then removed the neomycin resistant drug marker by crossing to a recombinase FLP derived mouse which recognizes the flippase recognition target (FRT) for mediated cleavage and generation of our founder mice. These mice were crossed to lineage-specific Cre-drivers to generate conditional knockout strains in liver (Albumin-Cre Jax #003574; (Postic et al., 1999)), neutrophils (S100a8-Cre Jax #021614; (Passegue et al., 2004)), and myeloid cells (LysM-Cre Jax #004781; (Clausen et al., 1999)). For all comparisons, Cre-positive and Cre-negative *Hmgcl^fl/fl^* littermates were used and genotypes were co-housed throughout lifespan until experimental endpoint. All experimental procedures were performed with approvals from the Yale and UCSF Institutional Animal Care and Use Committee.

### Physiological Measurements

For ketogenic diet experiments, mice were fed ad libitum ketogenic diet for up to one week, as indicated in each figure (89.5% of calories from fat, 10.4% of calories from protein, 0.1% of calories from carbohydrates; Research Diets D19042606). For obesity experiments, male mice were fed a high-fat diet (60% of calories from fat; Research Diets D12492) for 12 weeks. For fasting experiments, mice were fasted for 24hrs, beginning in the morning before tissue collection. For endotoxemia experiments, mice were challenged with LPS (O55:B5; 1mg LPS/kg body weight) and euthanized 4 hours later for analysis. Handheld meters were used to measure blood glucose (Contour Next) and BHB (Precision Xtra) levels in whole blood. For glucose tolerance tests, mice were fasted for 16 hours prior to intraperitoneal injection of glucose (0.6g glucose/kg body weight for old mice, 0.4g glucose/kg body weight). Lipolysis was assessed by measuring glycerol release from adipose tissue explants (Sigma #MAK117). Body temperature was measured with a rectal temperature probe (BAT-12 microprobe thermometer, Physitemp).

### Western blot

Neutrophils were isolated by magnetic negative selection (StemCell Technology) according to manufacturer’s protocol. For NLRP3 activation, cells were treated with LPS (1ug/ml, Sigma #L4391-1MG, strain 0111:B4) for 4 hours, followed by 45 minutes ATP (5mM, Sigma # A7699-1G). For all western blots, cell lysates were prepared in RIPA buffer containing protease and phosphatase inhibitors. Total protein concentrations were measured by BCA assay (Bio-Rad) and equal amounts of total protein were analyzed. Antibodies used were IL-1β (GeneTex # GTX74034), HMGCL (Proteintech # 16898-1-AP), Kbhb (PTM Biosciences #PTM-1201RM) and β-actin (Cell Signaling # 4967S).

### Flow Cytometry

Cells were made into single-cell suspension by filtering over 70μm filter. Red blood cells were lysed with ACK lysing buffer. Cells were incubated with Fc block and then stained with antibodies for standard lineage markers (CD3, B220, CD11b, Ly6G) for 30 minutes on ice, followed by three washes with FACS buffer, and then immediately acquired on a BD LSR II equipped with violet, red, green, and blue lasers, or an Attune NxT Flow Cytometer equipped with a blue, violet, green, and red laser. Data was analyzed with FlowJo (Ashland, OR). Antibodies were purchased from Biolegend.

### Gene Expression

mRNA was isolated from cells in QIAzol using the Qiagen RNeasy kit. cDNA was transcribed using the iScript cDNA synthesis kit (Bio-Rad). Gene expression was measured by RT-PCR by ΔΔC_t_ method and expressed relative to 18s (Table 1).

### Statistical Analyses

All graphs and statistical analyses were done in Prism (v9, GraphPad). For comparisons of 2 groups, two-sided student’s t-tests were used to calculate statistical differences. Comparisons of more than 2 groups were analyzed by 1-way ANOVA. To compare groups that were tracked over time, mice were individually tracked and statistical differences were calculated by paired 2-way ANOVA. P-values are provided in each figure.

## Acknowledgements

We thank all members of the Dixit and Goldberg labs for the thoughtful discussion and feedback about this project. The Goldberg lab is funded in part by 5R00AG058801, pilot awards from the UCSF NORC P30DK098722 and UCSF Liver Center P30DK026743, and the Chan Zuckerberg Biohub. The Dixit lab is funded in part by R01AR070811, P01AG051459, R01AG076782, R01AG068863, R01AG073969 and U54AG079759. We thank Yale Flow Cytometry for their assistance with flow cytometric service. The Core is supported in part by an NCI Cancer Center Support Grant # NIH P30 CA016359. We thank the UCSF Laboratory for Cell Analysis for access to flow cytometers. The LCA is funded by 5P30CA082103-23.

